# Novel uAUG creating variants in the 5’UTR of ENG causing Hereditary Hemorrhagic Telangiectasia

**DOI:** 10.1101/2022.12.18.520932

**Authors:** Omar Soukarieh, Emmanuelle Tillet, Carole Proust, Charlène Dupont, Béatrice Jaspard-Vinassa, Florent Soubrier, Aurélie Goyenvalle, Mélanie Eyries, David-Alexandre Trégouët

## Abstract

Hereditary Hemorrhagic Telangiectasia (HHT) is a rare vascular disorder causing abnormal vessel formation and characterized by autosomal dominant transmission. The associated considerable variability in symptoms and clinical severity complicate the management of the disease. In clinical routine, 3 main genes, *ACVRL1* (also known as *ALK1*), *ENG* and *SMAD4* are screened for pathogenic variants at the origin of HHT. About 80% of HHT cases are caused by pathogenic coding variants in *ACVRL1* and *ENG*. However, at least 15% remain with no molecular explanations in the 3 main genes. We here report the identification of 2 never reported variants, c.-79C>T and c.-68G>A, in the 5’UTR of *ENG* in 2 unrelated HHT patients. These 2 variants are predicted to create upstream AUGs (uAUGs) in the 5’UTR, which are in frame with a stop codon located within the CoDing Sequence (CDS), thus generating Overlapping upstream Open reading frames (uoORFs). In other cases, uAUGs can lead to fully upstream ORFs (uORFs) when associated with stop codons located within the 5’UTR or to elongated CDS (eCDS) when in frame with the CDS and associated with the main stop codon.

In order to assess the pathogenicity of these *ENG* variants, we performed *in vitro* functional assays based on the expression of wild-type and mutant constructs in human cells. We found that these 5’UTR *ENG* variants were associated with a decrease of protein levels in HeLa and HUVEC cells. They were also associated with a decreased ability to activate BMP9-stimulated ALK1 receptor in a BMP-response element (BRE) assay. This assay is a mandatory element before providing a definitive molecular diagnosis and has been so far applied only on coding *ENG* variants. We applied the same experimental workflow on 3 additional uoORF-creating variants (c.-142A>T, c.-127C>T and c.-10C>T) and one eCDS-creating variant (c.-9G>A) in the 5’UTR of *ENG* previously reported in HHT patients. We found that all the analysed uoORF-creating variants alter endoglin levels and function. A comparison of our experimental results with patients’ clinical characteristics suggests that uoORF-creating variants leading to ENG protein levels ≤ 40% *in vitro* would be associated with HHT. Additional experiments relying on an artificial deletion in our mutated constructs show that created uAUGs predicted to create uoORFs are able to initiate the translation indicating that the associated effect is likely caused by an alteration of the translation mechanism.

Overall, we here identified two never reported 5’UTR *ENG* variations in HHT patients and shed new lights on the role of upstream ORFs on ENG regulation. Our findings contribute to the amelioration of molecular diagnosis in HHT.

## Introduction

Hereditary Hemorrhagic Telangiectasia (HHT), also known as Osler-Weber-Rendu syndrome, is a rare vascular disorder characterized by an autosomal dominant transmission. It causes severe complications including bleeding, recurrent epistaxis, iron deficiency, anemia, mucocutaneous telangiectasias and arteriovenous malformations (AVM) in different organs (e.g. lungs, liver, brain, digestive tract). In some cases, AVM can be asymptomatic and hardly identified in HHT patients, thus risking stroke and cerebral abscess. In addition, there is a considerable inter- and intra-family variation in symptoms and clinical severity, even in cases resulting from an identical pathogenic variant ^1^. HHT is a multiorgan disease for which management and treatment depend on individual case evaluation and require a multidisciplinary approach. International guidelines have been elaborated for the diagnosis and management of HHT ^2,3^. According to these guidelines, anti-angiogenic and anti-fibrinolytic drugs are now recommended to treat bleeding and epistaxis in HHT patients ^3–6^ but treatment improvements are still awaited.

HHT affects about one in 5000 people ^1^ and is caused by pathogenic variants in genes coding for proteins of the Bone Morphogenetic Proteins (BMP) signaling pathway belonging to the transforming growth factor β (TGF-β) superfamily. For molecular diagnosis, 3 main genes, *ACVRL1, ENG* and *SMAD4* are routinely screened. It has been estimated that 80% of HHT cases are caused by pathogenic variants in *ENG* and *ACVRL1* genes encoding respectively endoglin and ALK1 proteins ^7,8^. Endoglin and ALK1 are part of a crucial signal pathway on endothelial cells. ALK1 is a serine-threonine kinase activating the phosphorylation of Smad1 and Smad5 in response to BMP9 or BMP10 ligands. Phosphorylated Smad1 and Smad5 will then bind to Smad4 and translocate into the nucleus to initiate specific gene transcription regulation. Endoglin has no kinase activity by itself but is able to enhance ALK1 signaling. A cellular-based assay has been developed to characterize coding missense variants in *ACVRL1* and *ENG* ^9,10^ but has never been used for non-coding variants. The developed assay allows evaluating the potential effect of coding *ACVRL1* or *ENG* variants on the ALK1 response to BMP9 by using a transfected reporter gene (luciferase) driven by a BMP-response element (BRE). This assay has been used to discriminate pathogenic from non-pathogenic variants and is now considered as a mandatory element in the molecular diagnosis of HHT. At least 15% of HHT cases remain without molecular explanations in exonic/flanking intronic regions of the 3 main causative genes ^11^ leading to diagnosis uncertainty for patients and their families. The remaining cases could be explained either by pathogenic variants in hypothetical undiscovered genes ^12–14^, or non-coding variants in known HHT genes. Indeed, the interpretation of non-coding variants, including promoter, 5’UTR, 3’UTR, and deep intronic variants, is a real challenge in molecular diagnosis and their characterization can contribute to resolve unexplained cases of HHT.

In the last 2 decades, a few studies have highlighted the pathogenicity of non-coding 5’UTR variants affecting upstream open reading frames (upORFs) which are important regulatory elements on the transcript ^15,16^. UpORFs result from the presence of an upstream Translation Initiation Site (uTIS) located within the 5’UTR and associated with an in-frame stop codon (uStop) located within the 5’UTR or the CoDing Sequence (CDS). Depending on the location of the uStop on the transcripts, one can dissociate different types of upORFs ^17^: (i) fully upstream ORF (uORF) when the uStop is located within the 5’UTR; (ii) overlapping upORF (uoORF) when the uTIS is out-of-frame with the CDS and is associated with a uStop codon located within the CDS; and (iii) elongated CDS (eCDS) when the upORF terminates at the main stop codon of the CDS. UpORFs can alter the protein levels by disturbing the translation initiation step and the recognition of the main TIS by the ribosomes. Of note, several upORF-altering variants have been shown to alter the protein levels and be pathogenic in humans. However, only few uAUG-creating variants have been classified deleterious in HGMD (Human Gene Mutation Database) and ClinVar databases ^17,18^, suggesting that such type of variants is underestimated.

Interestingly, non-coding variants creating uAUGs in the 5’UTR of ENG have been identified in HHT patients and reported in the literature ^19–22^ and in databases (https://www.ncbi.nlm.nih.gov/clinvar/?gr=1; https://www.hgmd.cf.ac.uk/ac/all.php; https://arup.utah.edu/database/ENG/ENG_display.php). Three of these variants (c.-142A>T; c.-127C>T and c.-10C>T) are predicted to generate uoORFs all ending at the same stop codon located at position c.125 in the CDS. Two of these variants (c.-142A>T and c.-127C>T) have been analyzed *in vitro* in published studies and have been associated with a decrease of endoglin levels ^19–21^. While c.-127C>T and c.-10C>T have been reported as pathogenic in public databases, the c.-142A>T is absent from databases but suggested to be pathogenic in Ruiz-Llorente *et al*., 2019 ^19^. An additional uAUG-creating variant (c.-9G>A) has been reported in the 5’UTR of *ENG* ^20^. As the created uAUG is in frame with the CDS, it is predicted to generate an eCDS. This variant has been associated with a very mild decrease of the endoglin levels *in vitro* ^20^ and its clinical interpretation is considered as “conflicting” or “pathogenic mild” in HGMD and HHT databases, respectively.

In this study, we focused on unresolved cases of molecular diagnosis of HHT. By reanalysing sequencing data of targeted genes from HHT patients, we identified 2 never reported variants in the 5’UTR of ENG predicted to create uAUGs in 2 unrelated patients. More precisely, these variants (c.-79C>T; and c.-68G>A) are predicted to create uoORFs also ending at the same stop codon located at position c.125 within the CDS. Using different functional *in vitro* assays, including original ones, we demonstrated that these variants alter endoglin levels and function, thus arguing in favour of a pathogenic effect of these variants. These assays were also applied to previously reported 5’UTR *ENG* variants.

## Materials and methods

### Clinical data of HHT patients and variant identification

Clinical diagnosis of HHT is determined based on the Curaçao criteria established by the HHT international committee ^23^. These criteria include the presence of epistaxis (spontaneous and recurrent nose bleeds), telangiectasias (multiple, at characteristic sites such as lips, oral cavity, fingers and nose), visceral vascular lesions (gastrointestinal telangiectasias and/or arterio-venous malformations), and family history (a first-degree relative with HHT). The diagnosis of HHT is robust if three of the criteria are present, possible or suspected if two are present and unlikely if fewer than two are present.

Genomic DNA was extracted from the whole blood of 274 individuals with suspected HHT collected at the genetics department of the Pitié-Salpêtrière Hospital (Paris, France). The patients provided written informed consent for their DNA material to be used for genetic analysis in the context of molecular diagnosis in accordance with the French bioethics’ laws (Commission Nationale de l’Informatique et des Libertés n° 1774128).

Genetic testing was performed by using a custom next-generation sequencing (NGS) targeted gene panel including HHT genes (*ACVRL1, ENG* and *SMAD4*, between others) and additional genes related to other hereditary vascular diseases. We applied the MORFEE bioinformatics tool ^24^ on vcf files from sequenced individuals in order to detect and annotate non coding SNVs creating uAUGs (uAUG-SNVs) in the 5’UTR of the sequenced genes.

### Plasmid constructs and expression in human cells

In order to evaluate the potential effect of *ENG* variants on the protein steady state levels, we performed the *in vitro* functional assay described by Labrouche et al. ^25^. To do so, we started by the amplification of the long isoform of ENG (L-ENG; NM_001114753.3) cDNA from HeLa cells by using specific primers covering the entire 5’UTR and the CDS lacking its stop codon (ENG c.-303_c.1974), and cloned it in the pcDNA3.1/myc-His(-) plasmid (Invitrogen) in frame with a Myc-His tag to obtain the wild-type (WT) clone (Supp. Table 1). The PCR reaction was performed using Phusion High Fidelity (HF) DNA polymerase (ThermoFisher) and the cloning was carried out after double digestion of the inserts and plasmid with BamHI and HindIII. Mutated clones carrying *ENG* variants identified in this study or described in the literature (Figure 1A, Table 1) ^19–22^ were prepared by directed mutagenesis on the WT generated clone, pcDNA3.1-L-ENG-WT (Figure 1B), using specific back to back primers (Supp. Table 1) and the Phusion HF DNA polymerase, followed by dpnI digestion of the matrix, phosphorylation of the generated PCR product, ligation and transformation of competent cells to obtain unique clones. In addition, we generated a supplemental construct in which we deleted the start codon of ENG (pcDNA3.1-L-ENG-c.1_2del, Supp. Table 1) and introduced separately *ENG* variants (c.-142A>T, c.-127C>T, c.-79C>T, c.-68G>A and c.-10C>T) in the latter construct. In presence of *ENG* c.1_2del, the created uAUGs become in frame with the Myc tag, allowing the detection of potential proteins translated from the uAUGs. All the recombinant plasmids were verified by Sanger sequencing (Supp. Table 1) (Genewiz).

**Table 1.**
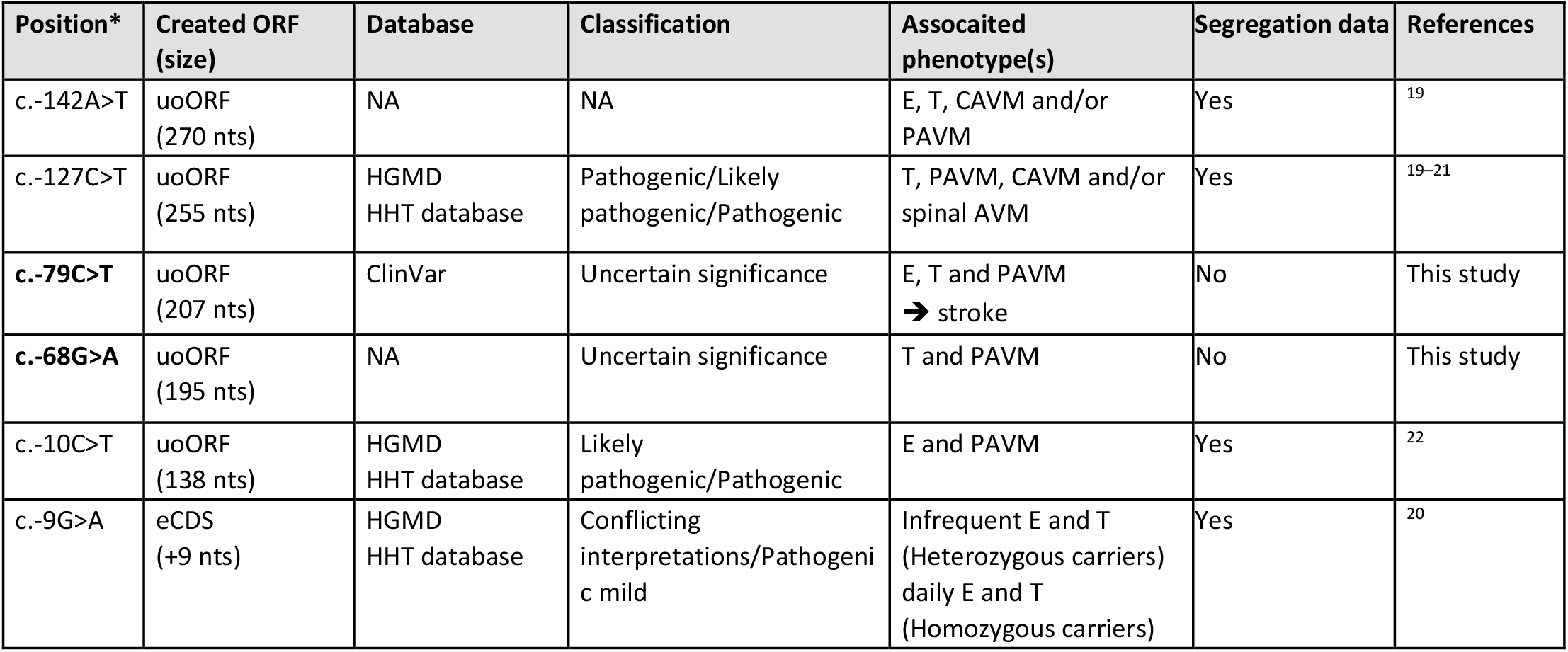
Bioinformatics predictions and clinical data for *ENG* 5’UTR variants analyzed in this study. *, variant position in the L-ENG NM_001114753.3 tanscript (c.1 corresponds to the A of the main AUG); uoORF, Overlapping upstream Open Reading Frame; eCDS, elongated CoDing Sequence; nts, nucleotides; HHT, Hereditary Hemorrhagic Telangiectasia; E, epistaxis; T, telangiectasia; NA, non-applicable; AVM, arteriovenous malformation; CAVM, cerebral AVM; PAVM, pulmonary AVM.

**Figure 1.**
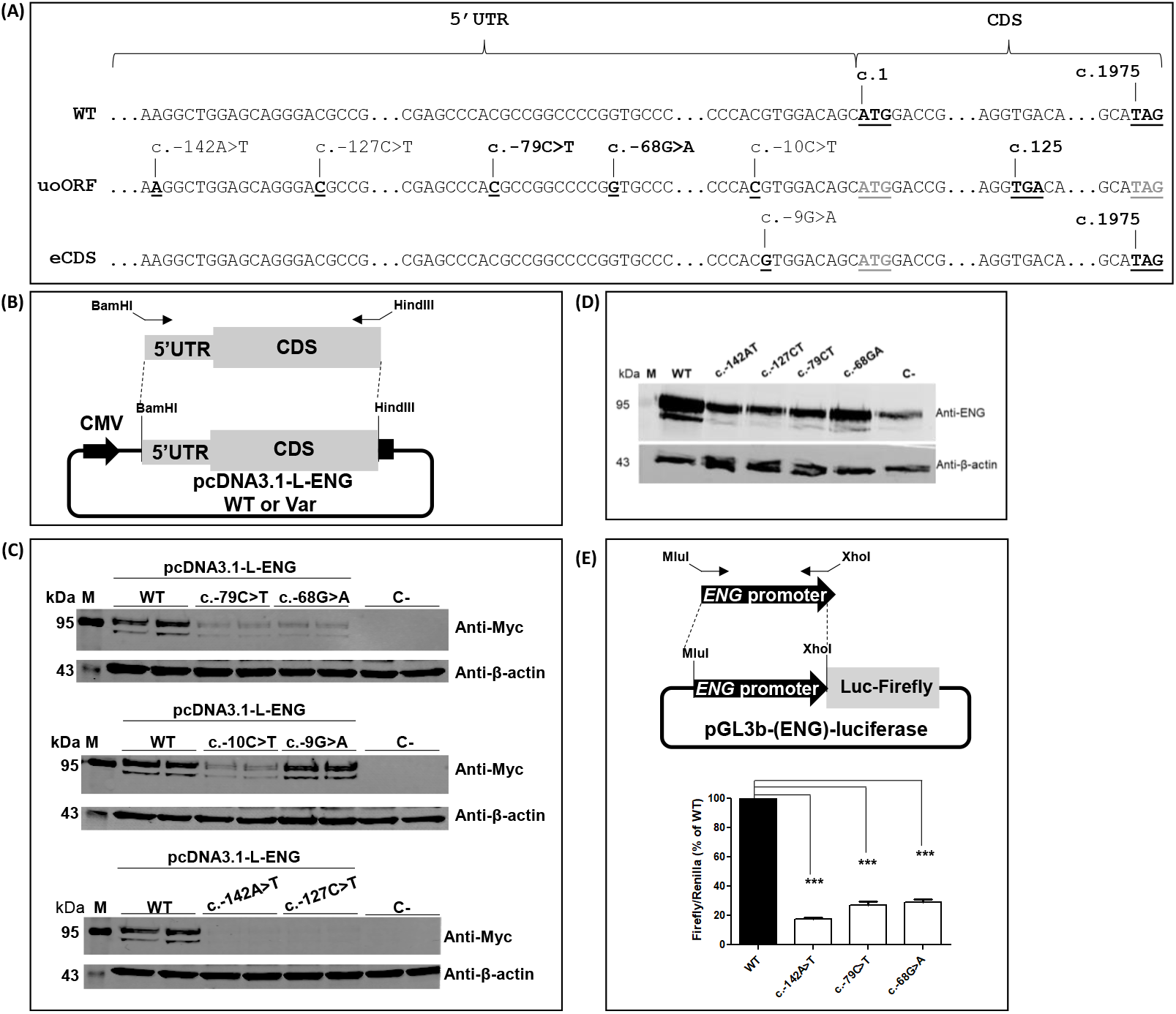
uAUG-creating variants are associated with a decrease of ENG levels *in vitro*. (A) cDNA of the long (NM_001114753.3) isoform of ENG transcripts. Positions of the identified 5’UTR uAUG-creating variants from this project and published studies ^19–22^ as well as position of the associated uStop codon at position c.125, and of main start (c.1) and stop (c.1975) codons are indicated. 5’UTR, 5’ Untranslated Region; CDS, CoDing Sequence; WT, wild-type; uoORF, overlapping upstream Open Reading Frame; eCDS, elongated CDS. (B) Schematic presentation of the pcDNA3.1-L-ENG constructs prepared and used for the evaluation of ENG steady-state levels in HeLa cells. Arrows indicate specific primers targeting the cDNA of ENG with extra sequences containing restriction sites to allow specific cloning in the expression vector. Position of the Myc-His tag is represented by the black square on the plasmid. CMV, cytomegalovirus promoter; WT, wild-type; Var, Variant; 5’UTR, 5’ UnTranslated Region; L, Long. (C and D) Western blot results on total proteins extracted from transfected HeLa cells with 1 µg of pcDNA3.1-L-ENG constructs or from transduced HUVEC cells with 20 MOI of lentiviruses containing ENG, respectively. Two bands of different molecular weights are observed for endoglin likely corresponding to more glycosylated (upper band) and less/non glycosylated (lower band) ENG monomers ^10^. Anti-Myc and anti-ENG correspond to the used antibodies for the target protein from HeLa and HUVECs, respectively, and anti-βactin corresponds to the antibody used against the reference protein. kDa, kilodalton; M, protein ladder; WT, wild-type, C-, negative control corresponding to pcDNA3.1-empty vector. Shown results are representative of 5 independent experiments. (E) Decrease of luciferase activity observed with uAUG-creating variants in *ENG*. Schematic presentation of the pGL3b-(ENG)-luciferase prepared and used in this assay is shown in the upper panel. Arrows indicate specific primers targeting the promoter of *ENG* with extra sequences containing restriction sites to allow specific cloning in the expression vector. Luc, luciferase. In the lower panel, shown results correspond to Firefly/Renilla ratios normalized to the wild-type (WT) in 5 independent experiments. ***, p-value < 10^−3^ (two-factor ANOVA followed by Tukey’s multiple comparison test of variants versus WT).

In order to evaluate the effect of the identified variants on protein levels, HeLa cells were transfected with the prepared pcDNA3.1-L-ENG constructs. HeLa were prepared in 6-well plates 24 hours before the transfection with 4.5.10^5^ cells/well in RPMI medium (Gibco-Invitrogen) supplemented with 10% fetal calf serum (Gibco-Invitrogen). Transfections were performed in duplicate with 1 µg of each plasmid using JetPRIME® reagent (Polyplus Transfection) according to the manufacturer’s recommendations. Empty pcDNA3.1/myc-His(-) plasmid was used as negative control. The day of the transfection, cell confluence was at 60-80%. Cells were harvested and lysed 48 hours after transfection to extract total RNA and protein. For this purpose, cells have been scraped and collected in 500 µl of PBS. Then, they were split in 2 aliquots: 100 µl aliquot for RNA extraction and 400 µl aliquot for protein extraction, as indicated below.

As endoglin is mainly known for its endothelial function in HHT, we also assessed the ENG steady-state levels in HUVEC cells (Human Umbilical Vein Endothelial Cells). For this purpose, we first used the generated pcDNA3.1-L-ENG constructs to subclone the WT and mutant cDNA of ENG in pRRLsin-MND-MCS-WPRE plasmid, upstream of a MND promoter. Unlike for pcDNA3.1-L-ENG constructs, pRRLsin-MND-MCS-WPRE-L-ENG ones do not contain any tag and the CDS of ENG ends at its own stop codon (ENG c.-303_c.1977). Then, lentiviruses were produced at the platform of vectorology (Vect’UB) of the University of Bordeaux (https://www.tbmcore.u-bordeaux.fr/vectorologie/) for the WT and *ENG* c.-142A>T, c.-127C>T, c.-79C>T and c.-68G>A variants. Twenty-four hours before transduction, HUVECs were prepared in 6-well plates with 2.8.10^5^ cells/well in EGM-2 medium (Endothelial Cell Growth Medium-2, Lonza). Cells were transduced in duplicate with 20 MOI (Multiplicity of Infection) of the generated lentiviruses by using 3.2 mg of protamine sulfate. Transduced cells were harvested with trypsin 72 hours post-transduction and each well was transferred to a P100 plate. For each construct, one plate has been used for whole protein extraction, 48 hours after the transfer to P100 plates and the duplicate plate was used to freeze transduced cells.

### Protein preparation and western blot analysis

Whole protein extractions were performed with RIPA supplemented with protease inhibitors for a pellet collected from one well of HeLa cells or one P100 plate of HUVECs. Concentrations were measured by using the BCA protein assay kit (Pierce™) following the manufacturer’s instructions. Proteins were loaded on 10% SDS-PAGE gels in parallel with a protein prestained ladder (Euromedex) and transferred onto PVDF membranes (Bio-Rad) by using the trans-blot turbo transfer system (Bio-Rad). Membranes were incubated with monoclonal anti-(c-Myc Tag) antibody (Merck Millipore) (HeLa extracts), anti-ENG (Abcam) (HUVECs extracts) to probe endoglin, and anti-β-actin (Cell Signaling) (HeLa and HUVECs extracts) as a loading control. Fluorescent goat anti-mouse IgG Alexa Fluor 700 (ThermoFisher) was used against anti-(c-Myc Tag) and goat anti-Rabbit IgG (H+L) Alexa Fluor 750 (Invitrogen) was used against anti-ENG and anti-β-actin. Odyssey Infrared Imaging System (Li-Cor Biosciences) in 700 and 750 channels was used to scan, reveal, and quantify the blots. For quantification, the average of each duplicate was computed from the quantified values and ENG levels for each sample were normalized to the corresponding β-actin levels then to the WT or the negative control levels (%). The two bands obtained for the Endoglin, corresponding to the more glycosylated (upper band) and less/non glycosylated (lower band) ENG monomers ^10^, were taken together for the quantification.

### RNA isolation and RT-qPCR analysis

In order to evaluate ENG transcript levels in transfected HeLa cells, total RNA was isolated from the collected pellets by using the RNeasy mini kit (Qiagen) following the manufacturer’s instructions. Extracted RNA was quantified and equal quantities were used for reverse transcription reaction, performed with the M-MLV reverse transcriptase (Promega). Then, 20 ng of cDNA of each sample was used for the qPCR reaction (duplicate/sample) with ENG or α-tubulin specific primers (Supp. Table 1) and the GoTaq qPCR Master mix (Promega), in presence of CXR reference dye, in a final volume of 10 µl and 40 cycles of amplification on QuantStudio3 Real-Time PCR System (ThermoFisher). QuantStudio design and analysis software was used to analyze the results and transcript levels were normalized to the reference α-tubulin gene. 2-ΔΔCT method was used to calculate the relative amounts of ENG to α-Tubulin in different samples. ΔΔCT were calculated by taking into account the mean of qPCR duplicates followed by the mean of transfection duplicates for each sample. Reaction efficiency (90%-110%) and melting curves were evaluated for each couple of primers.

### Luciferase assay

A complementary *in vitro* assay was deployed to evaluate the effect of the identified variants on the promoter activity. For this purpose, the *ENG* promoter, containing the basal promoter and the region carrying major transcriptional regulatory elements, as defined in Rıus et al., ^26^ was amplified by using specific primers (Supp. Table 1). This promoter sequence corresponds to the 805 nucleotides located upstream of the main ATG. The amplified promoter has been cloned in pGL3-basic vector containing the CDS of Firefly luciferase to obtain the WT clone (Figure 1E). Then, *ENG* variants c.-142C>T, c.-79C>T and c.-68G>A were introduced in parallel by directed mutagenesis, as described above for pcDNA3.1-L-ENG vectors. All the recombinant plasmids were verified by Sanger sequencing by using specific primers (Supp. Table 1) (Genewiz). WT or mutant clones were co-transfected with a plasmid containing Renilla luciferase in triplicate in 96-well plate of HeLa cells. Forty-eight hours after the transfection, luciferase activity was measured by using the dual-glo luciferase assay system (Promega) directly in transfected wells by detecting luminescence with both Firefly and Renilla luciferases. Mean of the triplicates of Firefly/Renilla ratios of each sample was normalized to the WT.

### Functional effect of *ENG* variants on BRE activity *in vitro*

The BRE assay described in Mallet et al. (11) has been modified to assess the functionality of the 5’UTR *ENG* variants. Briefly, NIH-3T3 were seeded in 96-wells white plates (15 000 cells/well) in DMEM containing 1% Fetal calf serum and transfected the following day by a mixture of plasmid (i) BRE luciferase reporter plasmid (75ng) (ii) pRL-TK luc encoding Renilla luciferase (20 ng), pcDNA3-ALK1 (0.15 ng) and pcDNA3.1-L-ENG WT or 5’UTR-mutated constructs (0 to 10 ng/well). Four hours after transfection, cells were stimulated overnight with 5 pg/ml of BMP9 in serum-free medium (R&D Systems) and luciferase activity was measured with the twinlite Firefly and Renilla Luciferase Reporter Gene Assay System (PerkinElmer). Means of triplicate were calculated for each sample and Firefly/Renilla ratio of stimulated wells was normalized to that obtained in non-stimulated wells.

### Statistical data analysis

Differential protein and RNA levels, luciferase activity, and BRE activity were assessed using two-factor ANOVA followed by Tukey’s multiple comparison test. A statistical threshold of p < 0.05 was used to declare statistical significance.

## Results

### uoORF-creating *ENG* variants are associated with a decrease of protein levels *in vitro*

We identified 2 variants located in the 5’UTR of *ENG* in 2 unrelated individuals with unresolved molecular diagnosis for HHT (Table 1). Both variants, c.-79C>T and c.-68G>A, are predicted to create uAUGs that are out-of-frame with the CDS and predicted to create uoORFs ending at the same stop codon located within the CDS (Figure 1A). The presence of these variants has been confirmed by Sanger sequencing. The individual carrying *ENG* c.-79C>T was clinically diagnosed with HHT since she presented epistaxis, mucosal and cutaneous telangiectasias and a pulmonary arteriovenous malformation that led to stroke. The individual carrying the *ENG* c.-68G>A had several telangiectasias, and pulmonary arteriovenous malformation.

These variants represent relevant candidates to explain HHT in these individuals but their pathogenicity still needed to be demonstrated. Apart from these 2 variants, other 5’UTR variants have been reported in *ENG*. Among them, 4 are at the origin of new uAUGs, which generate either uoORFs ending at the same stop codon in the CDS (c.-142A>T, c.-127C>T and c.-10C>T) or an elongated CDS (c.-9G>A) since the predicted uAUG is in frame with the CDS (Figure 1A, Table 1) ^19–22^. In order to contribute to the classification of the identified variants, we started by evaluating their potential effect on *in vitro* endoglin levels, in parallel with the published variants indicated above (Figures 1A-D).

First, we observed a decrease of the protein steady-state levels associated with both c.-79C>T and c.-68G>A variants in transfected HeLa cells, reduced to 17% and 42% relative to the WT, respectively (Figure 1C, Supp. Figure 1A). Interestingly, the published variant c.-10C>T was also associated with a decrease of endoglin levels in our assay (ENG levels ∼32%; Figure 1C, Supp. Figure 1A). Moreover, we observed a drastic effect (< 10% of WT levels) associated with c.-142A>T and c.-127C>T variants and a very moderate effect (∼80% of WT levels) observed with the c.-9G>A. RT-qPCR results on RNAs extracted from transfected cells showed similar levels of ENG between WT and variants (Supp. Figure 1B).

Similar results were obtained after overexpression of *ENG* variants in HUVEC cells. One can assume that protein levels obtained with the empty vector (C-on Figure 1D) reflect the endogenous levels of endoglin and are not disturbed by the transduction. Thus, western blot analysis on total proteins extracted from transduced cells revealed drastic effects for c.-142A>T and c.-127C>T variants with similar levels to those of the endogenous endoglin (Figure 1D, Supp. Figure 1C-D), followed by c.-79C>T with slightly higher levels (around 40% of WT levels), and then c.-68G>A associated with less pronounced effect (around 60% of WT levels).

Finally, we assessed the effect of *ENG* variants in the context of the promoter in a luciferase assay. We used this assay in order to study the potential effect of *ENG* variants on luciferase activity by altering the promoter activity and/or the translational mechanism in HeLa cells ^26,27^. We observed a decrease of the luciferase activity with the tested variants (c.-142A>T; c.-79C>T and c.-68G>A; Figure 1E). Importantly, the levels of luciferase appeared correlated to the protein levels detected for the same variants in the previous assays. At least for these variants, one could assume that the obtained effect on luciferase activity is related to the predicted uoORFs, also observed in the context of the pGL3b-(ENG)-luciferase construct, probably without any additional effect on the promoter activity. Again, these results show the alteration of ENG levels by the analyzed variants, in concordance with the two previous assays.

### Decreased levels of ENG related to 5’UTR variants alter the ability of ENG to activate BMP9-stimulated ALK1 receptor

The ability of *ENG* variants (Table 1) to enhance ALK1-mediated BMP9 response was assessed using a BRE assay in NIH-3T3 cells. First, we demonstrated that pcDNA3.1-L-ENG-WT construct used in this study had similar stimulation efficiency than the WT ENG clone used as a reference in previous studies (Supp. Figure 2A). We then tested 6 different 5’UTR variants and showed that the 5’UTR variants had a reduced ability to stimulate ALK1 (Figure 2). More precisely, c.-142A>T and c.-127C>T variants were associated with the lowest BRE activity detected (≤ 20% of WT levels), followed by variants c.-79C>T, c.-68G>A and c.-10C>T which induced a moderate activity (∼50% of WT levels) and finally variant c.-9G>A which was associated with a modest decrease of activity (∼70% of WT levels) (Figure 2). Of note, BRE activity obtained with this last variant was very variable between independent experiments, leading to conflicting interpretations of the alteration of activity by this variant. Altogether, the decrease of BRE activity obtained with the *ENG* 5’UTR variants correlated with the ENG levels measured *in vitro*. Moreover, BRE stimulation by *ENG* variants was hampered only when low levels of expression plasmid were transfected (below or equal to 1ng/well) (Supp. Figure 2B), highlighting that the lack of activity is directly linked to low ENG levels in the cells. Thus, these results demonstrate that uoORF-creating variants could be considered as hypomorphic variants.

**Figure 2.**
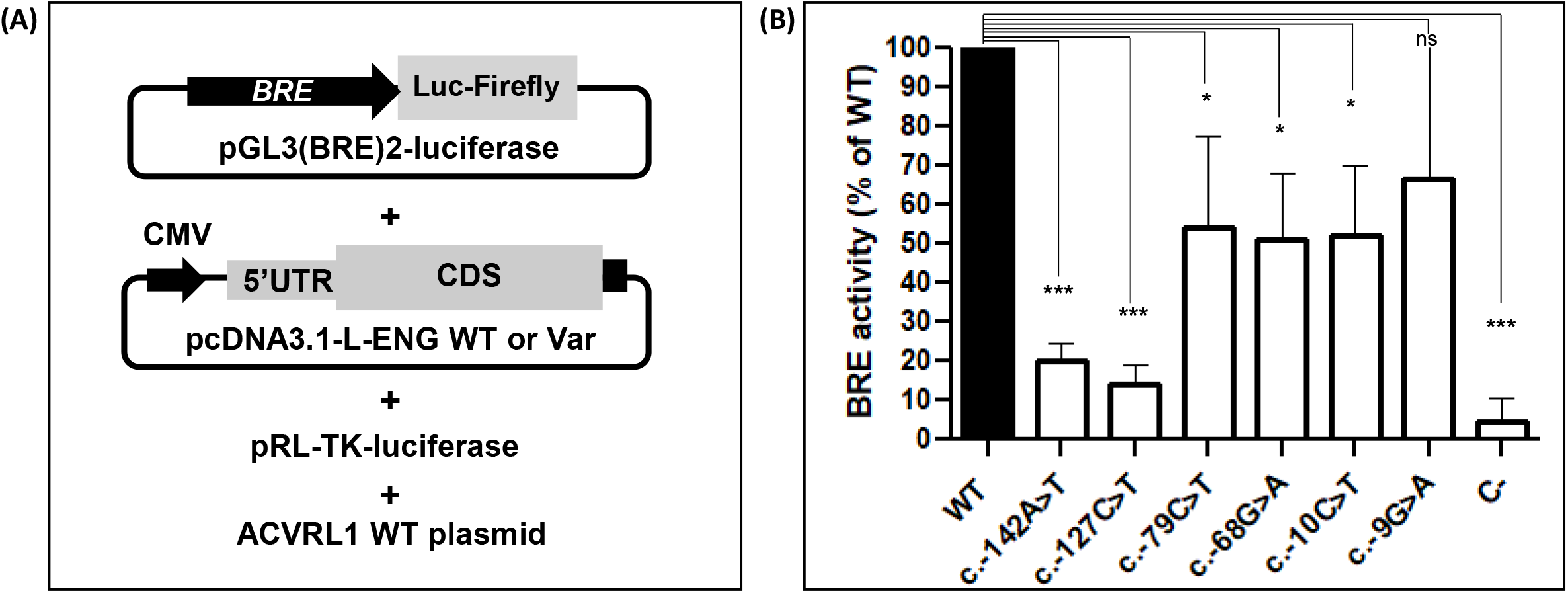
ALK1 response to BMP9 stimulation is affected by 5’UTR *ENG* variants. (A) Schematic presentation of the co-transfected constructs in this assay. BRE, BMP-response element; CMV, cytomegalovirus promoter; WT, wild-type; Var, variant; 5’UTR, 5’ UnTranslated Region; L, Long. (B) Decrease of BRE activity observed with all the analyzed variants in co-transfected NIH3T3 cells stimulated with BMP9 (5 pg/ml). Shown results correspond to the quantification of Firefly/Renilla and normalized to the wild-type (WT) (n = 4). ***, p-value < 10^−3^; *, p-value < 5.10^−2^, ns, non-significant (two-factor ANOVA followed by Tukey’s multiple comparison test of variants versus WT).

### The created uAUG in the 5’UTR of ENG are able to initiate the translation

We next investigated whether the uAUGs created by *ENG* variants in the 5’UTR could be used to initiate the translation of a new protein. In order to answer this question, we started by suppressing the main ATG of *ENG* by deleting the dinucleotide *ENG* c.1_2del (Figure 3). This deletion is predicted to completely abolish the translation of the CDS. In addition, in the presence of any of the *ENG* uAUG-creating variants, this deletion should transform the uoORFs into an elongated CDS as the created uAUGs become in frame with the main stop codon. That leads to identify potentially translated proteins from uAUGs *in vitro*. pcDNA3.1-L-ENG carrying the deletion with or without 5’UTR variants were transfected in HeLa cells and ENG levels were assessed in western blot and RT-qPCR as described above.

**Figure 3.**
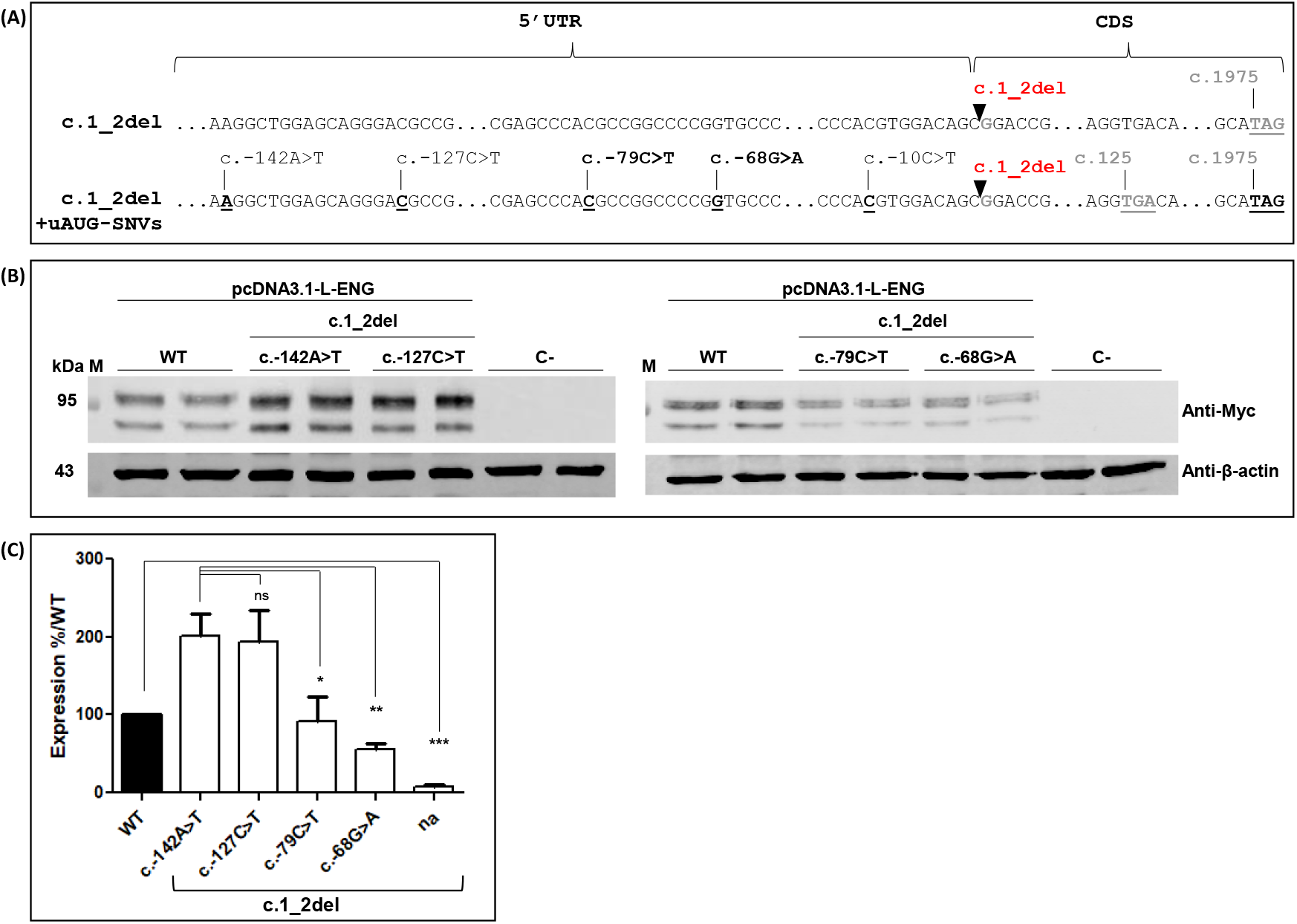
Created uAUGs in the 5’UTR of ENG seem to be able to initiate the translation. (A) cDNA of the long (NM_001114753.3) isoform of ENG transcripts. Positions of the identified 5’UTR uAUG-creating variants from this project and published studies ^19–22^ as well as position of the associated uStop codon (c.125), the introduced deletion c.1_2del, and of main start (c.1) and stop (c.1975) codons are indicated. 5’UTR, 5’ Untranslated Region; CDS, Coding sequence; WT, wild-type; uoORF, overlapping upstream Open Reading Frame. (B) Western blot results on total proteins extracted from transfected HeLa cells with 1 µg of pcDNA3.1-L-ENG constructs. Two bands of different molecular weights are observed for endoglin likely corresponding to more glycosylated (upper band) and less/non glycosylated (lower band) ENG monomers ^10^. Anti-Myc and anti-βactin correspond to the used antibodies for the target and the reference proteins, respectively. kDa, kilodalton; M, protein ladder; WT, wild-type, C-, negative control corresponding to pcDNA3.1-empty vector. Shown results are representative of 5 independent experiments. (C) Quantification of protein steady-state levels obtained in (B) and probably resulting from translation initiation at predicted uAUGs. For quantification, the average of each duplicate has been calculated from the quantified values and ENG levels for each sample have been normalized to the corresponding β-actin levels then to the WT (%). na, no 5’UTR variant introduced. Graphs are representative of 5 independent experiments. ***, p-value < 10-^3^; **, p-value < 10-^2^; *, p-value < 5.10-^2^, ns, non-significant (two-factor ANOVA followed by Tukey’s multiple comparison test of variants versus WT).

Interestingly, we detected high protein levels associated to c.-142A>T and c.-127C>T variants in the presence of the deletion (> 190% in comparison to the WT construct) and less important levels with c.-79C>T and c.-68G>A (∼90% and 50%, respectively) (Figures 3B-C). Similar results were also obtained for c.-10C>T (Supp. Figures 3A-B). The observed proteins appear to be of similar size to the WT product. Considering that the predicted size difference between the predicted elongated proteins initiated at the different uAUG and the WT protein is only 0.4 to 4.8 kDa, they would be undistinguishable in our Western blots. Importantly, these levels are inversely proportional to the levels of endoglin detected in our *in vitro* assays in association with the studied variants. Indeed, the highest protein levels obtained in this test were associated with variants that had the most drastic effect on ENG levels *in vitro* (Figures 1 and 3). These results would suggest that the obtained effects with *ENG* variants are related to a potential competition between created uAUGs and the main AUG at the translation level. One could notice that a very low amount of proteins (< 10% of the WT levels) is also detected with the pcDNA3.1-L-ENG-c.1_2del construct that could result from the use a non-canonical translation initiation site in the 5’UTR of ENG (Figure 3C, Supp. Figure 3A). Apart for c.-68G>A, relative ENG transcript levels were similar in presence of the deletion with or without uAUG-creating variants as detected by RT-qPCR (Supp. Figure 3C). Combined effect of these variants (at least for the c.-68G>A) on the transcription and/or RNA stability cannot be excluded.

## Discussion

The starting point of this project was the identification of 2 new variants in the 5’UTR of ENG (c.-79C>T and c.-68G>A) in 2 independent patients diagnosed with HHT but with unresolved molecular diagnosis. Here, we evaluated the effect of these variants on endoglin levels and function using *in vitro* assays. Data accumulated in this study suggest that these two variants are responsible of a decrease in endoglin levels (17-42% of WT levels in HeLa cells). These results were reproduced in HUVECs, thus showing the relevance of the *in vitro* assay in HeLa cells. Moreover, we found that these variants are associated with a decrease of BRE activity *in vitro* (50% of the WT). Mallet et al. have previously considered that missense *ENG* variants associated with partial response to BMP9 (40-60%) were pathogenic. Indeed, they showed that variants they studied with partial response are associated with confirmed diagnosis of HHT and three of them showed familial co-segregation ^10^. Based on this study, one can consider that the two hypomorphic variants we identified are likely responsible of the HHT phenotypes in these patients. In addition, we here evaluated for the first time the functional effects of *ENG* c.-10C>T variant. We show that this variant is associated with ∼32% of protein steady-state levels in HeLa cells and with ∼50% of BRE response. This variant is classified as likely pathogenic in HGMD based on family history and patient’s phenotype ^22^. As a consequence, data accumulated in this work suggest classifying c.-79C>T and c.-68G>A variants also as likely pathogenic. Furthermore, our results obtained for *ENG* c.-142A>T that parallel those we observed for *ENG* c.-127C>T, classified as pathogenic in HHT database, and combined with published data ^19^ add strong support for this variant being pathogenic. The observed effects are concordant with those obtained by other groups ^19–21^. However slight difference in the quantified endoglin levels could be observed and may be due to differences in the used constructs and cells ^19–21^.

While the two uoORFs we identified and three out of the four published upORFs start at different uAUGs located within the 5’UTR, they all end at the same stop codon located at position c.125. The only exception holds for c.-9G>A variant, that creates a uAUG in frame with the CDS, and generates an elongated CDS, probably at the origin of a longer form of the ENG protein carrying 3 additional amino acids. In this study, we assessed the potential functional effect of all of these variants. Curiously, a weak decrease (∼20%) of ENG levels and BRE activity have been associated with c.-9G>A variant compared to a drastic reduction observed for c.-142A>T and c.-127C>T and moderate reduction for c.-79C>T, c.-68G>A and c.-10C>T. Interestingly, uoORF-creating variants (Figure 1A, Table 1) are all associated with confirmed HHT diagnosis associated with severe symptoms. Concerning the 3 published uoORF-creating variants, molecular findings are consistent with clinical and familial data, suggesting that uoORF-creating variants in *ENG* can cause a severe form of HHT. This hypothesis still needs to be confirmed by analysing all additional uoORF-creating variants in *ENG*, even those associated with non-uAUG codons. Of note, all uoORF-creating variants studied in this report and identified in severe forms of HHT were associated with ENG protein levels ≤ 40% *in vitro* (HeLa cells) and with ≤50% of BRE activity. Additional analysis would be interesting in order to define thresholds of pathogenicity of *ENG* variants in these assays. Furthermore, additional data will be mandatory to clarify the classification of c.-9G>A eCDS-creating variant.

Cellular-based BRE activity assay has been used to characterize *ENG* missense variants identified in HHT patients. Here, we adapted this assay to study 5’UTR variants in *ENG* and we observed that this kind of variants could alter the response of ENG to BMP9. However, while missense variants seem to alter the expression of ENG at the membrane and/or have negative effect on endogenous ENG, 5’UTR variants more likely cause a decrease of ENG levels in cells, which then results in an impaired response to BMP9. Thus, one could suggest that uAUG-creating variants associated with a decrease of ENG levels *in vitro* will probably alter the BRE activity and be likely pathogenic.

In total, three complementary assays have been used in this project to evaluate the functional effects of *ENG* 5’UTR variants and they all provided concordant results, suggesting that they could be reliably used for the functional characterization of upORF-variants in *ENG*.

Again, we demonstrated that uAUG-creating variants in *ENG* could alter the protein levels and function. However, this could happen via different mechanisms ^28,29^. Indeed, upORFs are part of the most known translational regulatory elements. They could, for example, enter in competition with the main coding sequence and affect the translation of this latter. In order to assess the potential translation of the created uoORFs, we started by deleting the main ATG, in the presence of uAUG-creating variants. Our results suggest that the created uAUGs could initiate the translation. More precisely, by deleting 2 nucleotides, we transformed overlapping upORFs into elongated CDS. That led us to identify proteins that are probably initiated at the uAUGs. We combined these data with bioinformatics predictions to estimate the translation confidence of the created uAUGs in the 5’UTR of ENG by using the PreTIS tool based on the combination of 44 features calculated from mRNA sequence (https://service.bioinformatik.uni-saarland.de/pretis/) ^30^. We applied PreTIS predictions on the 6 variants and obtained 0.67 (Low) to 0.96 (High) scores of translation confidence for the created uAUGs (Supp. Table 2). At least for the variants evaluated in this study, translation could be initiated at uAUGs carrying PreTIS scores ≥ 0.67 in ENG. However, these scores do not seem to be able to predict the strength of the translation initiation and still need to be evaluated. Moreover, we analysed the Kozak sequence surrounding the created uAUGs and found that those resulting from the c.-142A>T and c.-127C>T variants are surrounded with stronger kozak sequences comparing to the 3 other uoORF-creating variants (c.-79C>T, c.-68G>A and c.10C>T) (Supp. Table 2). These observations are consistent with our results obtained *in vitro* showing a higher amount of protein associated with the c.-142A>T and c.-127C>T variants (Figure 3). The precise identification of the detected proteins will require further investigations using supplemental methods. In addition, it would be interesting to assess the functional potential of these proteins in the BRE activity assay. Indeed, if these proteins show some restoration of ENG functions, this may lead to new therapeutic approaches common to all uoORF-creating variants ending at the same uSTOP. At last, Kim and collaborators showed a decrease of RNA levels in carriers of the c.-127C>T variant ^21^ and c.-142A>T has been predicted to create a binding site for the transcription regulatory factor HOXA3 ^19^, suggesting that potential effect of the identified variants on the transcription cannot be excluded.

Finally, *ENG* is one of the rare examples of genes that are rich in uAUG-creating variants in the 5’UTR. Our study, between others, demonstrated that 5’UTR variants predicted to create upORFs should not be neglected in molecular diagnosis of genetic diseases. While we only studied uAUG-creating variants in the 5’UTR of *ENG* here, we are aware that 5’UTR variants creating non canonical translation initiation codons or disrupting existing upstream ORFs, should also be given more attention.

