## Supplemental Figures and Tables for "Novel uAUG creating variants in the 5’UTR of ENG causing Hereditary Hemorrhagic Telangiectasia"

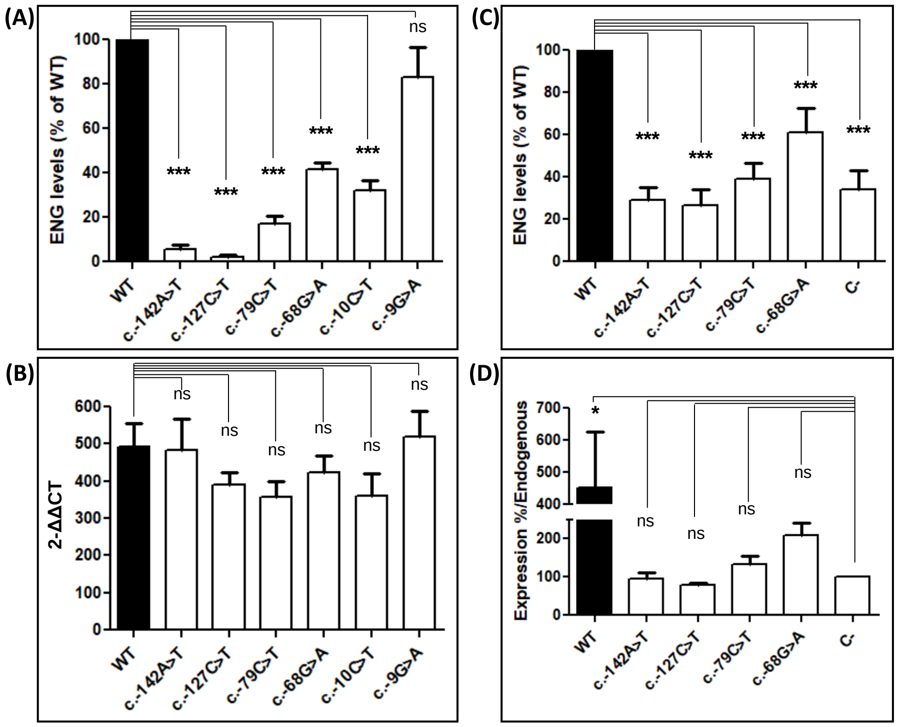


**Supplemental Figure 1. Quantification of ENG levels in HeLa (A-B) and HUVEC cells (C-D).** (A) Apart for c.-9G>A variant, ENG steady-state levels in HeLa cells is significantly decreased with variants in comparison to the wild-type (WT) construct. For quantification, the average of each duplicate has been calculated from the quantified values and ENG levels for each sample have been normalized to the corresponding β-actin levels then to the WT (%).The two bands obtained for the Endoglin, corresponding to the more glycosylated (upper band) and less/non glycosylated (lower band) ENG monomers (1), were taken together for the quantification. (B) RNA levels of Endoglin do not variate between wild-type and variants. RT-qPCR results are shown. 2-ΔΔCT corresponds to the normalization of ENG to internal control α-tubulin, then to negative control (empty vector). (C) Decrease of ENG protein levels in HUVECs cells in comparison to the wild-type (WT). ENG levels for each sample have been normalized to the corresponding β-actin levels then to the WT (%). C-, Negative control corresponding to the empty vector. (D) ENG levels associated to variants are similar or slightly higher of endogenous endoglin (C- on the Figure). ENG levels for each sample have been normalized to the corresponding β-actin levels then to the negative control (C-) (%).The graphs are representative of 5 independent experiments. ***, p-value < $10^{-3}$, *, p < $5.10^{-2}$, ns, non-significant (two-factor ANOVA followed by Tukey's multiple comparison test of variants versus WT (A, B, C) or negative control (D)).

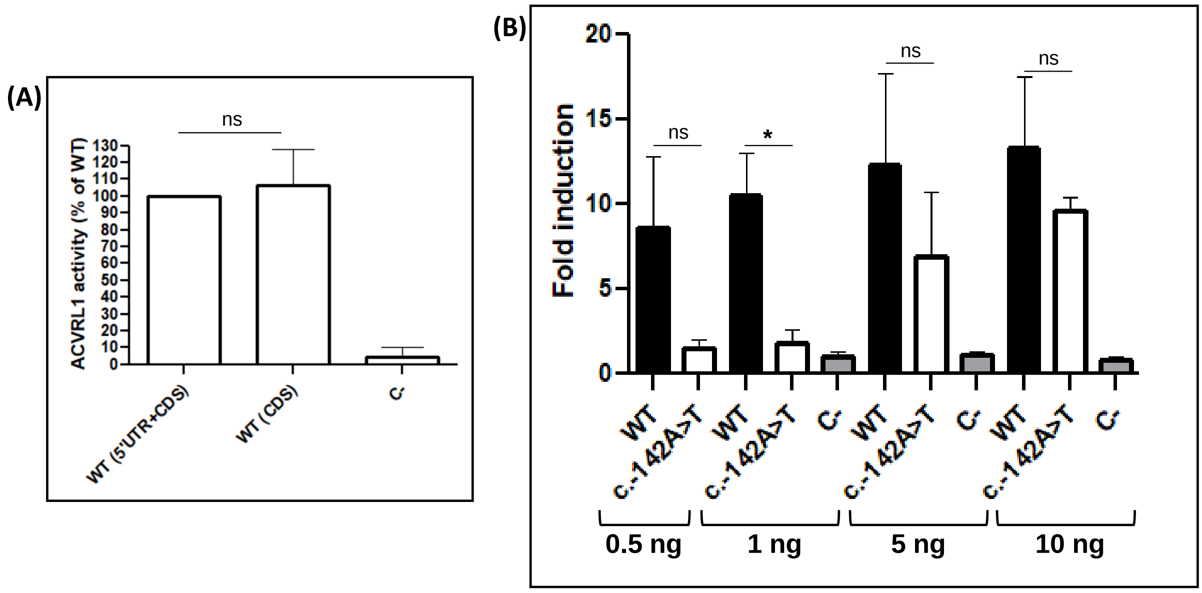


**Supplemental Figure 2. Optimization of cellular-based BRE activity assay to be applicable on *ENG* 5'UTR variants.** (A) pcDNA3.1-L-ENG-WT vector gives similar BRE activity to ENG-WT plasmid only containing the CDS of ENG (WT(CDS)) used in Mallet et al., 2015 (1). C- corresponds to pcDNA3.1 empty vector used as negative control. WT, wild-type; 5'UTR, 5' UnTranslated Region; CDS, CoDing Sequence. The graph is representative of 4 independent experiments. ns, non-significant (two-factor ANOVA followed by Tukey's multiple comparison test of WT(5'UTR+CDS) versus WT(CDS)). (B) Titration of ENG vectors with increasing quantities in order to determine optimal conditions to study 5'UTR variants. Fold induction corresponds to the detected activity normalized to control without ENG (C-). Transfected quantities of pcDNA3.1-L-ENG-WT or variant plasmids, and empty vector (C-) in 96 well plates are indicated. The graph is representative of 3 independent experiments. ng, nanograms; ns, non-significant; *, $p < 5.10^{-2}$ (two-factor ANOVA followed by Tukey's multiple comparison test of c.-142A>T variant versus WT).

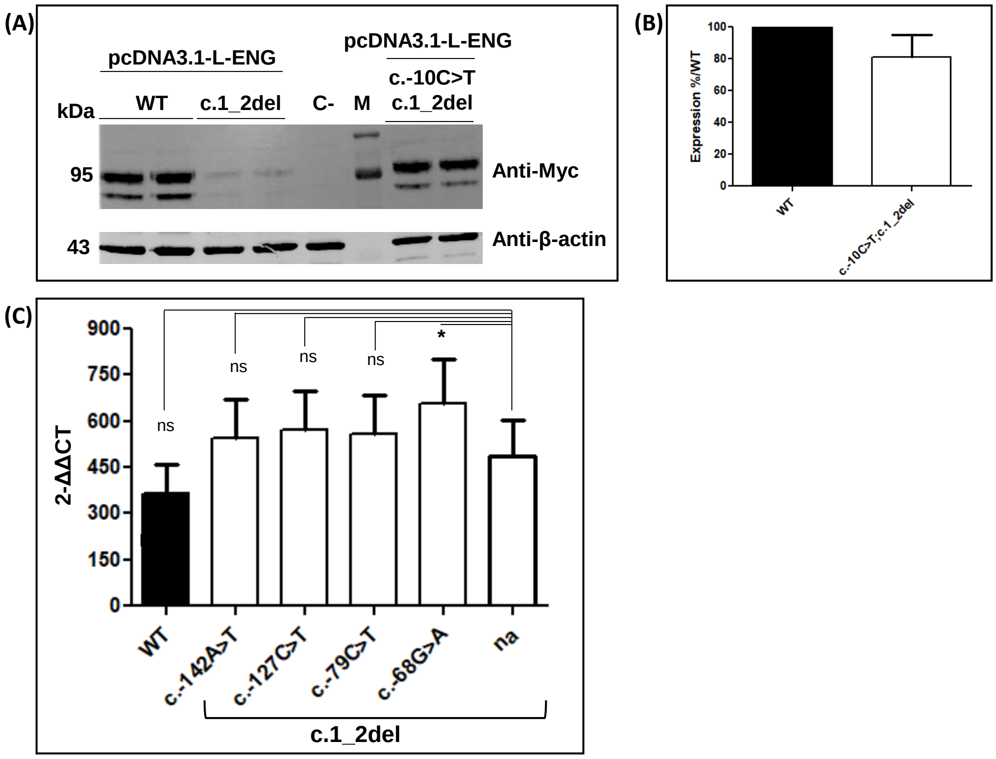


**Supplemental Figure 3. Created uAUGs in the 5'UTR of ENG seem to be able to initiate the translation.** (A) Western blot results on total proteins extracted from transfected HeLa cells with 1 µg of pcDNA3.1-L-ENG constructs. Two bands of different molecular weights are observed for the Endoglin likely corresponding to more glycosylated (upper band) and less/non glycosylated (lower band) ENG monomers (1). Anti-Myc and anti-βactin correspond to the used antibodies for the target and the reference proteins, respectively. kDa, kilodalton; M, protein ladder; WT, wild-type, C-, negative control corresponding to pcDNA3.1- empty vector. (B) Quantification of protein steady-state levels obtained in (A) and probably resulting from translation initiation at the created uAUG in presence of the c.-10C>T variant. For quantification, the average of each duplicate has been calculated from the quantified values and ENG levels have been normalized to the corresponding β-actin levels then to the WT (%). Graphs are representative of 5 independent experiments. *, p-value < $5.10^{-2}$, ns, non-significant (two-factor ANOVA followed by Tukey's multiple comparison test of variants versus WT). (C) Quantification of ENG RNA levels in HeLa cells transfected with pcDNA3.1-L-ENG constructs containing the c.1_2del deletion in presence or in absence (na) of uAUG-creating variants. RT-qPCR results are shown. 2-ΔΔCT corresponds to the normalization of ENG to internal control α-tubulin, then to negative control. Graphs are representative of 5 independent experiments.

**Supplemental Table 1. Primers used in this study for cloning, qPCR, back-to-back directed mutagenesis (DM) and sequencing.** Restriction sites are underlined. Variations are in lowercase. Position of c.1_2del is indicated with a dash.

| Technique | Plasmid | Primer name | Primer sequence (5'-3') |
|---|---|---|---|
| Cloning | pcDNA3.1 & MND14 | ENG-Iso1-WT-BamHI-F | tagtggatccATACCACAGCCTTCATCTGCGC |
| Cloning | pcDNA3.1 | ENG-Iso1-WT-HindIII-R | gcccaagcttTGCCATGCTGCTGGTGGAG |
| Cloning | pRRLsin-MND-MCS-WPRE | ENG-Iso1-NheI-R | gcaggctagcCTATGCCATGCTGCTGGTGGAG |
| Cloning | pGL3b | ENG-Prom-MluI-F | tcttacgcgtGGATCCCAGCGCTACCATCTTC |
| Cloning | pGL3b | ENG-Prom-XhoI-R | agatctcgagGCTGTCCACGTGGGGGC |
| qPCR | Human ENG | ENG-qPCR-Ex9-F | ACAAGTTTGTCTTGCGCAGT |
| qPCR | Human ENG | ENG-qPCR-Ex11-R | CTGTCCATGTTGAGGCAGTG |
| qPCR | Human α-Tubulin | hu_TUBA F | GATGCTGCCAATAACTATGCCCGAG |
| qPCR | Human α-Tubulin | hu_TUBA R | GAAAACCAAGAAGCCCTGAAGACGG |
| DM | pcDNA3.1-ENG | ENG_c.-142AT-F | CACCCCAGAtGGCTGGAGC |
| DM | pcDNA3.1-ENG | ENG_c.-142AT-pR | GCGGCCGAGGGGTCAG |
| DM | pcDNA3.1-ENG | ENG_c.-127CT-F | GAGCAGGGAtGCCGTCGC |
| DM | pcDNA3.1-ENG | ENG_127CT-pR | CAGCCTTCTGGGGTGGC |
| DM | pcDNA3.1-ENG | ENG-c.-79CT-F | GAGCCCAtGCCGGCC |
| DM | pcDNA3.1-ENG | ENG-c.-79CT-pR | GCACGGGGACCCGAGGGG |
| DM | pcDNA3.1-ENG | ENG-c.-68GA-F | GGCCCCGaTGCCCGC |
| DM | pcDNA3.1-ENG | ENG-c.-68GA-pR | GGCGTGGGCTCGCACGGG |
| DM | pcDNA3.1-ENG | ENG-c.-10CT-F | CAGGCCCCCAtGTGGACA |
| DM | pcDNA3.1-ENG | ENG-c.-9GA-F | CAGGCCCCCACaTGGACA |
| DM | pcDNA3.1-ENG | ENG-c.-10_-9-pR | TGCGCTGGGCCTTATCCTG |
| DM | pcDNA3.1-ENG | ENG-c.1_2delAT-F | CACGTGGACAGC-GGACCGCGGC |
| DM | pcDNA3.1-ENG | ENG-c.-2_-1delGC-pR | GGGGCCTGTGCGCTGG |
| Sequencing | Human ENG | ENG-seq1-F | CATGTCCTCTTCCTGGAGTTC |
| Sequencing | Human ENG | ENG-seq2-F | ATGTCCTTGATCCAGACAAAG |
| Sequencing | pcDNA3.1 | CMV-F | CGCAAATGGGCGGTAGGCGTG |
| Sequencing | pcDNA3.1 | BGH-R | TAGAAGGCACAGTCGAGG |
| Sequencing | pRRLsin-MND-MCS-WPRE | Fvf MND BclXl | ACCCTGTGCCTTATTTGAAC |
| Sequencing | pRRLsin-MND-MCS-WPRE | verifMNDhGli3 | GGCATTAAAGCAGCGTATCC |
| Sequencing | pRRLsin-MND-MCS-WPRE | R Arhgef11 MND | ATCCACATAGCGTAAAAGGAGCA |
| Sequencing | pGL3b | GL primer2 | CTTTATGTTTTTGGCGTCTTCCA |
| Sequencing | pGL3b | RV primer 3 | CTAGCAAAATAGGCTGTCCC |

**Supplemental Table 2. Bioinfomatics predictions and kozak sequences of the created uAUGs in ENG.** PreTIS translation confidence scores for created uAUGs are indicated. Optimal Kozak sequence surrounding an AUG as defined in Kozak, 1984 (2) was taken into account c**R**ccAUG**g** (R = purine, A or G) to compare the sequences surrounding the created uAUGs. Nucleotides at positions -3 and +4 relative to the predicted uAUGs are in bold (3,4). Position the uAUG-creating variant is underlined. The sequence surrounding the main AUG of ENG is c**a**gcATG**g.** *, variant position in the L-ENG; NM_001114753.3 transcript (c.1 corresponds to the A of the main AUG).

| Position* | PreTis Score (Interpretation) | Kozak Sequence |
|---|---|---|
| c.-142A>T | 0.84 (High) | c**c**agAAG**g** |
| c.-127C>T | 0.84 (High) | a**g**ggACG**c** |
| c.-79C>T | 0.67 (Low) | g**c**ccACG**c** |
| c.-68G>A | 0.89 (High) | c**c**cgGTG**c** |
| c.-10C>T | 0.85 (High) | c**c**ccACG**t** |
| c.-9G>A | 0.96 (High) | c**c**acGTG**g** |
